# Making Neurobots and Chimerical Ctenophores

**DOI:** 10.1101/2024.10.28.620631

**Authors:** Leonid L Moroz, Tigran P. Norekian

## Abstract

Making living machines using biological materials (cells, tissues, and organs) is one of the challenges in developmental biology and modern biomedicine. Constraints in regeneration potential and immune self-defense mechanisms limit the progress in the field. Here, we present unanticipated features related to self-recognition and ancestral neuro-immune architectures of new emerging reference species - ctenophores or comb jellies. These are descendants of the earliest survival metazoan lineage with unique tissues, organs and independent origins of major animal traits such as neurons, muscles, mesoderm, and through-gut. Thus, ctenophores convergently evolved complex organization, compared to bilaterians. Nevertheless, their neural and immune systems are likely functionally coupled, enabling designs and experimental construction of hybrid neural systems and even entire animals. This report illustrates impressive opportunities to build both chimeric animals and neurobots using ctenophores as models for bioengineering. The obtained neurobots and chimeric animals from three ctenophore species (*Bolinopsis, Mnemiopsis*, and *Pleurobrachia*) were able to be autonomous and survive for days. In sum, the unification of biodiversity, cell biology, and neuroscience opens unprecedented opportunities for experimental synthetic biology.

## Introduction

The origin of animals (Metazoa) has been a long-standing enigma ever since Darwin. Molecular reconstructions of ancestral toolkits are equally challenging due to the poor representation of basal metazoans as reference species. One of the critical puzzles is to decode the origin of multicellularity as the capability to resolve conflicts of individuality - the potential to recognize self (allorecognition) underlying clonal or aggregation mechanisms (Nicotra, 2019;Blackstone, 2021;Hiebert et al., 2021;Kapsetaki et al., 2023;Gui et al., 2024;Scott, 2024) and social interactions (Gherardi et al., 2012). Many organisms in complex ecosystems under environmental stress ‘come’ across the same fundamental dilemma: *To fuse or not to fuse* (Brusini et al., 2013).

Historically, in quests to decipher ancestral allorecognition toolkits, a lot of attention has been focused on sponges and cnidarians (Theodor, 1970;Powell et al., 2011;Gundlach and Watson, 2018;Rodriguez-Valbuena et al., 2022), which were perceived as the most “primitive” animal lineages. However, during the last decade, the animal tree of life has been under substantial revision, placing morphologically complex ctenophores, which previously were considered animals with bilaterian-grade organization, as the sister lineage to the rest of Metazoa (Ryan et al., 2013;Moroz et al., 2014;Whelan et al., 2015;Whelan et al., 2017;Schultz et al., 2023). Today, we view ctenophores (or comb jellies) as the survived descendants of the earliest branch on the animal Tree of Life with independently evolved canonical metazoan traits such as neurons, muscles, mesoderm, and through-gut (Moroz et al., 2014;Moroz, 2015;Moroz and Kohn, 2016).

There are 185 accepted ctenophore species (Moroz et al., 2024), and probably at least a hundred species await their discoveries. Most ctenophores are pelagic and fragile organisms with high regeneration potentials, recognized for more than a century (Chun, 1880;Mortensen, 1913;Tanaka, 1932;Coonfield, 1936;Coonfield, 1937a;Dawydoff, 1938;Freeman, 1967;Henry and Martindale, 2000).

B.R. Coonfield was a pioneer in comparative studies of self-recognition, and he performed more than a dozen different types of animal fusion experiments at the Marine Biological Laboratory (MBL, Woods Hole, MA, USA) using *Mnemiopsis leidyi* (Coonfield, 1937b) - the most abundant Atlantic ctenophore. Coonfield elegantly demonstrated virtually all possible combinations of grafting transplantations of different parts of the ctenophore body, including the surprising fusion of two or three animals together and the ability to generate animals with two aboral organs and complex integration (Coonfield, 1937b). Coonfield also stressed ‘*the power to regulate itself*’” in *Mnemiopsis*, and pointed out the coupling the animal symmetry and intrinsic mechanisms maintaining the aboral-oral axis (Coonfield, 1937b) or morphological homeostasis, in modern terms.

Some of Coonfield’s experiments were recently repeated (again at the MBL), confirming the capability of two individual *Mnemiopsis* to be fused following their injury and physical contacts (Jokura et al., 2024), with the rapid development of coordination between feeding lobe contractions within two hours and merging of meridional canals (a network of internal canals transporting nutrients, gametes as components of digestive and reproductive systems).

Inspired by Coondield’s systematic research (Coonfield, 1937b) and recent work on *Mnemiopsis* (Jokura et al., 2024), we asked questions about how broadly the fusion capabilities can occur across ctenophore species. Or whether only *Mnemiopsis* can be such a unique model.

Here, we show that at least two other ctenophore species (*Bolinopsis microptera* and *Pleurobrachia bachei*) are capable of fusion and rapid formation of chimeric animals. Furthermore, our surgical experiments show the capability of these two species and *Mnemiopsis* to form reduced organoid-type entities, which we named *ctenobots* and *neurobots*, a sort of simpler ‘living machines’, opening unprecedented opportunities for further exploration of emerging integrative properties, with insight into bioengineering, homeostatic intercellular signaling, and animal evolution.

## Results

*Bolinopsis microptera* (Moser, 1907) and *Pleurobrachia bachei* (A. Agassiz, in L. Agassiz, 1860) are two abundant species in the Pacific Northwest with remarkably different morphologies, lifestyles, and systematic positions (**Fig. 1**). *Pleurobrachia* represents a more basal branch of ctenophores with a canonical cydippid morphology and a pair of long tentacles for prey capture. In contrast, *Bolinopsis* and *Mnemiopsis* belong to a more derived clade Lobata, with characteristic wide feeding lobes in adults (although lobates preserve the cydippid stages earlier in their life cycles). Genomes of these species have been sequenced with 13 assembled chromosomes (Moroz et al., 2014;Hoencamp et al., 2021;Schultz et al., 2023;Koutsouveli et al., 2024). These cydippid and lobate ctenophores represent two distinct bodyplan organizations and feeding strategies.

**Figure 1.**
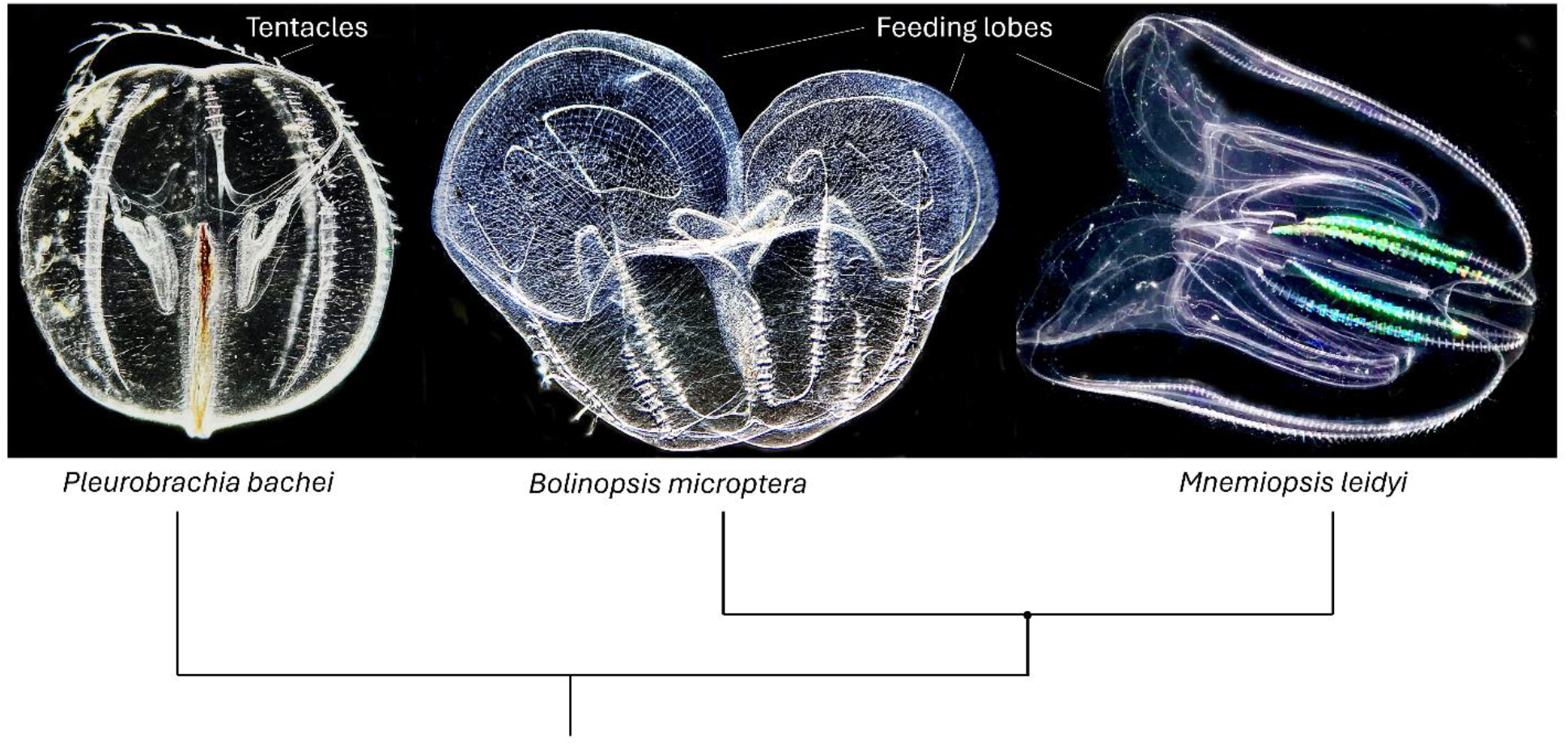
Three reference ctenophore species and their phylogenetic relationships (from (Moroz et al., 2014). *Pleurobrachia bachei* (A. Agassiz, in L. Agassiz, 1860) represents a more basal branch of ctenophores with a canonical cydippid morphology and a pair of long tentacles for prey capture (Yip, 1984;Townsend et al., 2020). *Bolinopsis* and *Mnemiopsis* (family Bolinopsidae) belong to Lobata, with characteristic feeding lobes in adults (MAIN, 1928;Colin et al., 2010;Granhag and Hosia, 2015;Cordeiro et al., 2022).

### Fusion of individuals in ctenophores

#### Fusion in Bolinopsis

Following the injury and removal of one feeding lobe and part of their body (exposing mesoglea) and placing two animals together with their injured sides facing each other for 12-20 hours (see Methods), we repeatedly formed fused chimeric *Bolinopsis* (N=11). In 12 hours, all wounds were healed, and the complete integration of two fused animals was evident (**Fig. 2**). The fused animals were maintained in a sea tank for one week, capable of locomotion and feeding.

**Figure 2.**
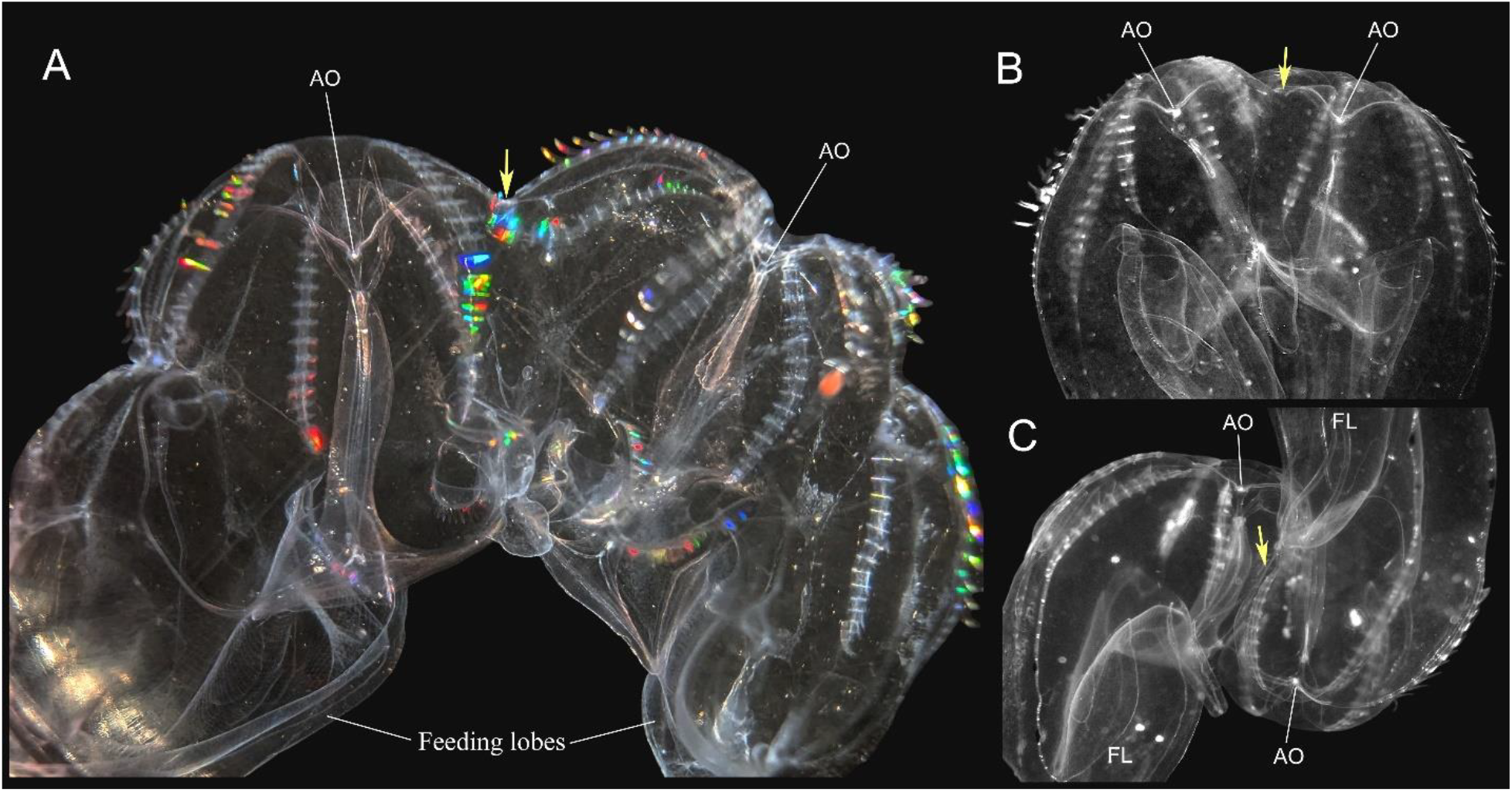
Fusion in *Bolinopsis microptera* (1 day). A – Two animals forming a chimera with far-spread aboral organs facing the same direction. All internal structures from both animals are visible. B - Chimera with closely positioned two aboral organs, which has body proportions similar to an intact single animal. Note the absence of any visual indication of the fusion boundary); C – Fused animals have been rotated 180 degrees and have a reverse (aboral-to-mouth) position. Yellow arrows indicate the sites of fusion. Abbreviations: AO – aboral organ; FL – feeding lobes.

There were three types of fusion configurations that we performed. In the first and main group, we preserved the same axis configuration placing animals side-by-side with aboral organs facing the same direction (**Fig. 2A,B**). Depending on how deep the cuts of the body wall were made, some chimeric animals were very wide with aboral organs far away from each other (**Fig. 2A**), while other animals had very closely positioned aboral organs and looked like a single animal of normal proportions but with two closely located aboral organs (**Fig. 2B**). In the second group of experiments, we rotated one animal 180° with a resulting chimera having aboral organs (and mouths) facing opposite directions (**Fig. 2C**). And in the third group of experiments, we dissected an aboral “cup” from one animal and replaced it in the second animal essentially grafting the aboral organ from one *Bolinopsis* to another.

The aboral region of ctenophores contains the aboral organ, the analog of ‘the elementary brain’ with the gravity sensor (statolith), and associated polar fields (Moroz, 2024). Thus, it is possible to graft the ‘elementary brain’ between two genetically different individuals.

*Bolinopsis* chimeras were visually healthy and actively swimming on days 1-to-7 Animals that were fused in the traditional orientation (aboral-to-aboral) showed coordination during swimming and looked like intact *Bolinopsis*. They could swim forward, reverse their direction to backward swimming, and take turns – further indicating coordinated behaviors. Their swimming was constant as in conrol individual animals. All fused *Bolinopsis* in response to a touch stimulus to one animal showed a robust and short-latency contraction response in both animals, mostly in their feeding lobes areas, as well as a brief inhibition of cilia beating (N=26). A similar response occurred when one of the animals touched the wall of an aquarium spontaneously during swimming – there was a strong contraction response in both animals (N=12).

Animals that were fused in the reversed position (aboral-to-mouth) were also swimming with active macrocilia beating. However, they did not move around as fast as the aboral-to-aboral chimeras, and their locomotion appeared to be less coordinated. However, they also showed a well-coordinated response to a tactile stimulus – a single touch of one feeding lobe caused a robust and short-latency (< 1 sec) contraction of lobes in both fused animals (N=9).

#### Fusion in Pleurobrachia

To our surprise, we also observed a reliable fusion in another ctenophore species, *Pleurobrachia* (N=8; **Fig. 3**), which is not known for such robust regenerative abilities as *Bolinopsis* and *Mnemiopsis*. The fusion occurred at a significantly slower rate by a factor of two. The complete epithelial wound healing was observed only on the second day. However, the extensive mesoglea fusion was obvious the next day, 12 hours after dissection.

**Figure 3.**
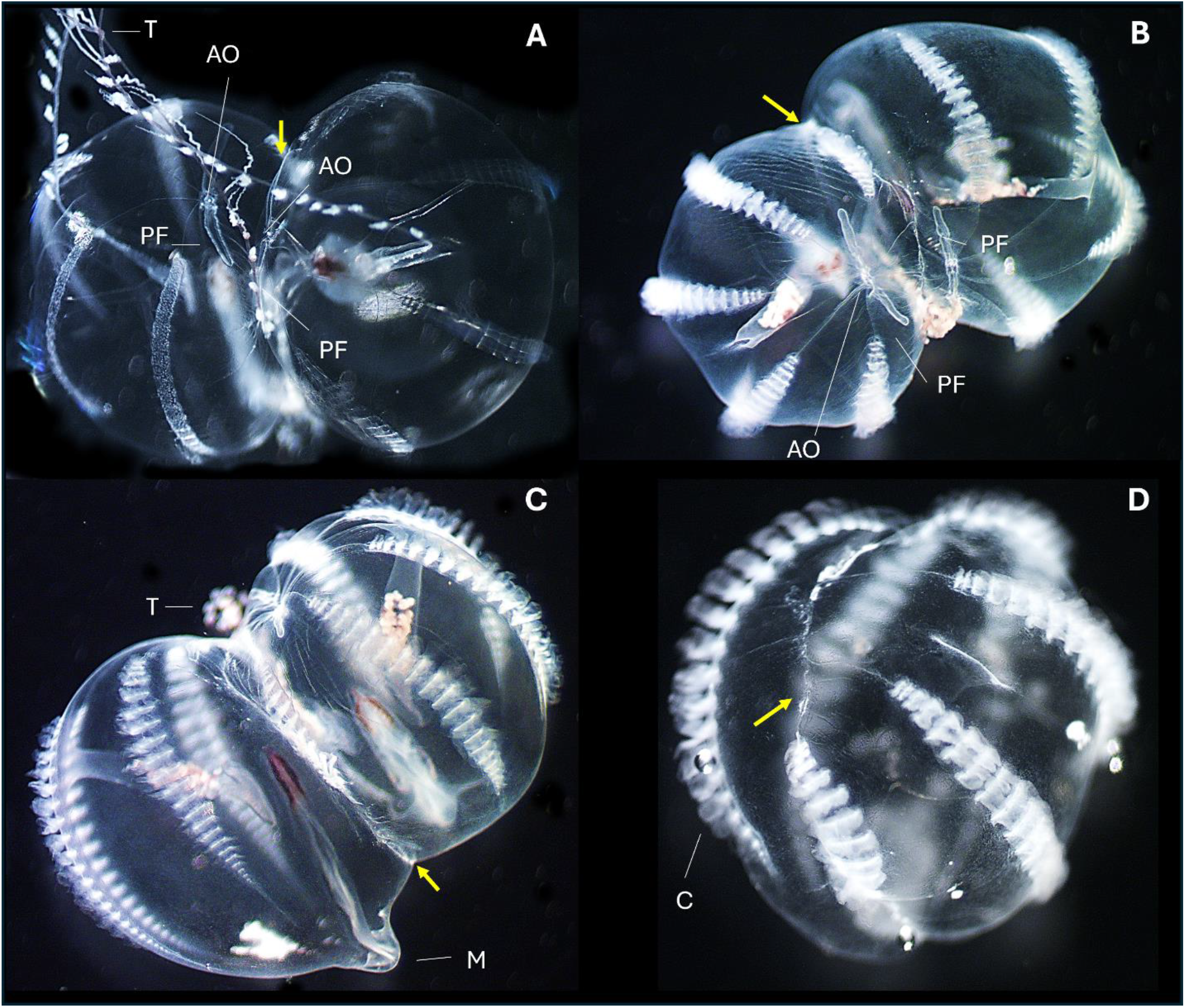
Fusion in *Pleurobrachia bachei* (2 days). A – Live chimera consisted of two fused animals, freely swimming in an aquarium; B-C – Fixed chimeras, with two aboral organs and two digestive tracts from different viewpoints. The yellow arrow indicates the fusion scar, which could be seen better after fixation. D – Small fused chimera without aboral organs or digestive tracts. The arrow indicates the fusion scar. Note that the orientation of ctene rows is not aligned between the parts from different animals. Abbreviations: AO – aboral organ; C – combs; M – mouth; PF – polar fields; T – tentacles.

We performed two types of fusion experiments with *Pleurobrachia*. In the first group, the animals retained their aboral organs and the resulting chimeras had two aboral organs and digestive tracts (**Fig. 3A-C**). In the second group, the leftover small parts of *Pleurobrachia* without aboral organs and digestive tracts were fused together, resulting in small chimeras lacking both (**Fig. 3D**). Both small (without aboral organs) and large (with two aboral organs) *Pleurobrachia* chimeras were very active moving around the container non-stop with constant swim cilia beating. In contrast to *Bolinopsis*, in *Pleurobrachia*, we did not see the regeneration of the polar bodies, or aboral organs or combs, when they were removed or injured.

Our behavioral experiments in *Bolinopsis* showing well-established coordination between two parts of the chimera suggested that their neural networks restore their connections and integrity after fusion. Although the fusion of *Pleurobrachia* was slower, it was equally reliable, offering better opportunities for microscopic analysis. In contrast to *Bolinopsis*, this species is better suited for fixation and follow-up mapping studies. It allows us to address one critical point of behavioral observations on fused animals – neuronal bases of coordination between two parts of the chimera and the possibility that neural networks restore their integrity after fusion in ctenophores.

Here, using the tubulin antibody immunolabeling for visualization of neural elements in ctenophores (Norekian and Moroz, 2016;2019), we checked whether the polygonal subepithelial neural networks from two chimera parts indeed establish connections between each other and unite the networks into one functional system. We received such confirmation, clearly identifying new neural processes that crossed the fusion scar and connected the neural networks from two previously independent animals (**Fig. 4**). Thus, the polygonal networks establish new connections that unite two fused parts.

**Figure 4.**
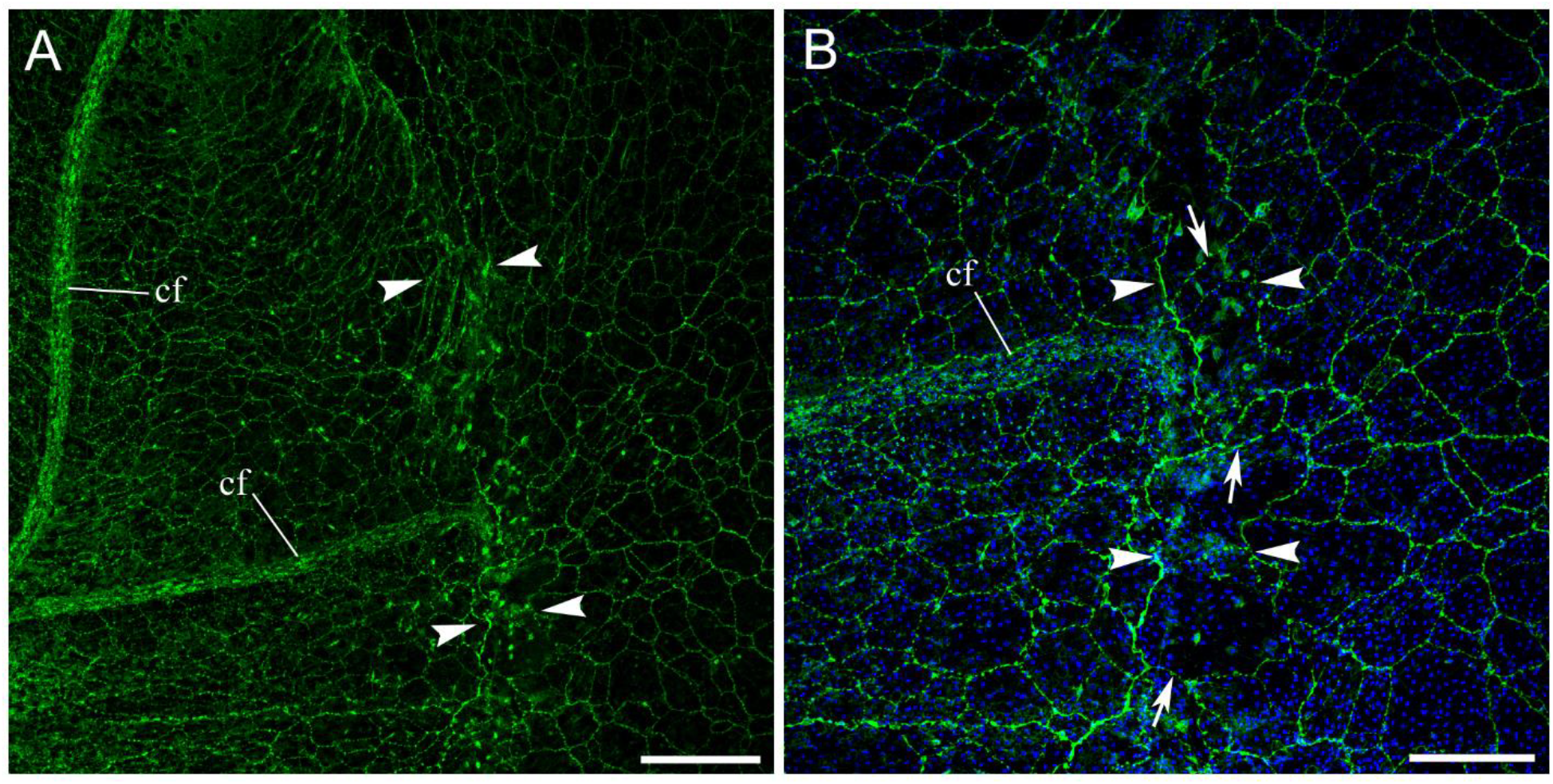
Continuity of subepithelial polygonal neural networks in *Pleurobrachia* chimera. A – polygonal subepithelial neural networks in two merged animals revealed by tubulin immunolabelling (in green). The fusion location is outlined by arrowheads. B – Higher magnification of the fusion area (nuclei are stained by DAPI in blue). Arrowheads outline the fusion scar area. Arrows point to the neural processes connecting the polygonal networks from both merged parts of *Pleurobrachia*. These images are from the 2^nd^ day of recovery. Abbreviation: *cf* – ciliated furrow. Scale bars: A - 200 µm; B - 100 µm.

### Making Ctenobots and Neurobots

By performing a variety of microsurgical procedures on *Bolinopsis* and *Mnemiopsis*, we noted that isolated small (submillimeter sizes) and larger (centimeter sizes) fragments from virtually any part of the ctenophore body (starting from the aboral organ to combs and auricles) can heal their wounds and close extensive injury sites within 1.5-6 hours, forming free organoid-like structures (**Fig. 5A-J**), which can be maintained as autonomous swimming entities for several days in seawater without adding nutrients.

**Figure 5.**
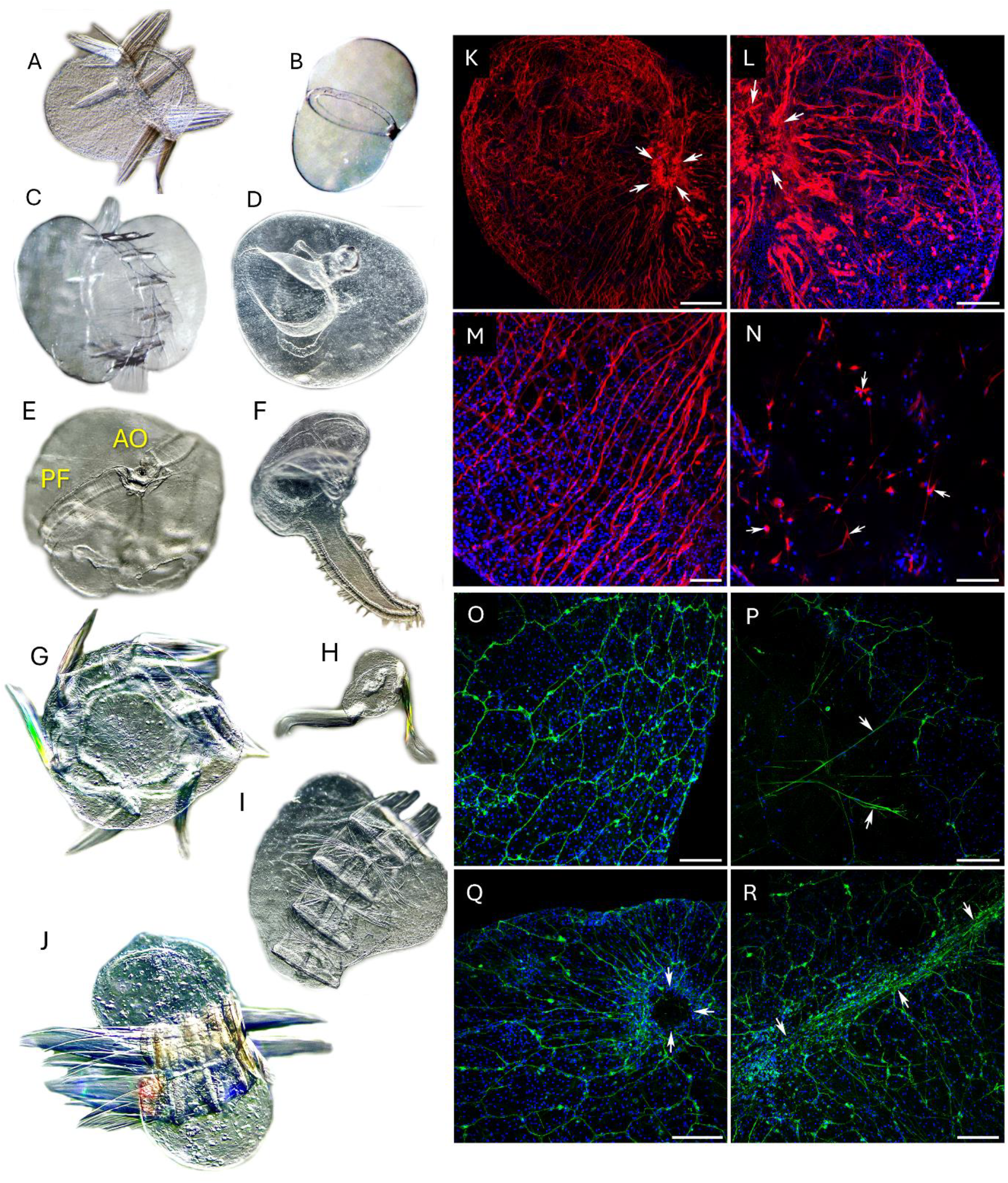
Neurobots and ctenobots. All images are from live preparations. A-J illustrate examples of different ctenobots from *Mnemiopsis leidyi* (A-C) - A and C contain reduced comb rows, while B possesses one internal meridional canal. D-J illustrate ctenobots from *Bolinopsis microptera*. D – mouth area and one meridional canal. E – isolated aboral region with the aboral organ (AO) and polar fields (PF); F – a ctenobot from the auriculus; G – five combs fused circularly. H – two fused reduced combs; I-J – four fused asymmetric combs with associated muscle bands. These entities were able to swim with combs and show muscle contractions.\*\*\**Bolinopsis* ctenobots labeled by phalloidin (in red) have an intact muscle structure and clearly identifiable fusion area (nuclei are stained by DAPI in blue). K – Small ctenobot 2-3 mm in diameter created from the body wall next to the feeding lobes. Arrows point to the fusion scar area. L, M – Higher magnification of different areas of the same ctenobot showing individual muscle fibers inside the tissue. Arrows point to the fusion scar. N – Mesogleal region inside the ctenobot. Arrows point to some of the mesogleal cells that are labeled by phalloidin. O-R: *Bolinopsis* ctenobots have polygonal subepithelial neural networks (labeled by tubulin antibody in green) as the intact adult animals do (nuclei are stained by DAPI in blue). O – Polygonal neural network. P – Mesogleal region in ctenobots contains long neural fibers (arrows). Q – The neural network is denser around the closing wound area (arrows point to the wound). R – The neural network is also very dense in the fusion scar area where two large pieces of tissue merged. Scale bars: K - 200 µm; L, O-R - 100 µm; M, N - 50 µm

We called these systems *ctenobots* or *neurobots* with the following distinction. Ctenobots we defined as self-maintained organoids with limited coordination and swimming capabilities, and these systems do not contain comb plates. Neurobots have a greater degree of autonomy, incorporate combs, and are capable of performing coordinated locomotory patterns with different degrees of oscillations and rhythmicity under apparent neuronal controls with reduced neural nets preserved and functional.

Both ctenobots and neurobots were able to contract spontaneously or following mechanical stimulation with preservation of mechanoreception and polarized muscles over 2-7 days of observations. Labeling of bots using phalloidin (**Fig. 5K-N**) revealed extensive meshwork of muscles and, we guess, potential mechanoreceptors (Norekian and Moroz, 2020;2021;2024b), which can explain their responses to mechanical stimulation and contractivity. Tubulin immunoreactivity also revealed the preservation of extensive canonical epithelial polygonal nerve nets and mesogleal neurons (**Fig. 5O-R**), similar to those in intact preparations for the same species (Norekian and Moroz, 2020;2021;2024a).

These types of neurobots were not formed from *Pleurobrachia* tissue, which can be explained by the fact that *Pleurobrachia* has less regeneration potential than lobates.

## Discussion

Reduced allorecognition and the ability to form chimeras are limited across the animal kingdom (Theodor, 1970;Rosengarten and Nicotra, 2011;Karadge et al., 2015;Chang et al., 2018;Melillo et al., 2018;Nicotra, 2019;Mueller and Rinkevich, 2020;Hiebert et al., 2021;Buckley and Dooley, 2022), and beyond (Gibbs et al., 2008;Kapsetaki et al., 2023), implying the exceptionality of this adaptation strategy. On the other hand, phenotypic plasticity in ctenophores (as descendants of the earliest branched metazoan lineage) can be significant. It is reflected in the astonishing diversity of forms across extant and extinct ctenophores (Moroz et al., 2024) as well as reverse development in *Mnemiopsis* (Soto-Angel and Burkhardt, 2024).

The ability to merge two individuals is a prominent characteristic of three currently investigated ctenophore genera: *Mnemiopsis* (Coonfield, 1937b;Jokura et al., 2024), *Bolinopsis*, and *Pleurobrachia* (this study). It further emphasizes the spectrum of ontogenetic plasticity in ctenophores. Notably, in the White Sea, Dr. Alexander Semenov observed *Beroe cucumis* with two mouths (Bezio et al., 2024). Because *Beroe* is nested within the Lobata clade (Moroz et al., 2014;Whelan et al., 2017), we might interpret this 2-mouth *Beroe* as either a result of abnormal development or a ‘fusion event’ similar to other lobates.

Our attempt to perform fusion experiments with another gelatinous pelagic animal from the same location was not successful. For example, we tested the similar-sized hydrozoan jellyfish *Clytia gregaria* (Order Leptomedusae, family Campanulariidae; this species was formerly known as *Phialidium gregarium*) with the same protocols and did not see any evidence of fusion (N=5).

Is reduced allorecognition in ctenophores a primary ancestral feature, or has this trait evolved independently? The current data on *Pleurobrachia*, together with two lobates, suggest that such capabilities were present in the common ancestor of the Pleurobrachiidae+Bolinopsidae clade. Still, it would be important to perform fusion experiments on more basal lineages, such as *Euplokamis, Mertensia*, and Platyctenida, to obtain a more detailed reconstruction of ancestral and derived features related to regeneration/allorecognition traits and associated mechanisms.

Mechanisms of the described ctenophore fusion are currently unknown. Initial genomic studies suggest the reduced complement of immunity-related genes in ctenophores (Moroz et al., 2014;Traylor-Knowles et al., 2019;Koutsouveli et al., 2024), lacking such canonical regulators as ikβ, NF-kβ-like, and MyD88 (Traylor-Knowles et al., 2019). However, alternative immune players are expected (Vandepas et al., 2024).

Comparative studies on regeneration in *Vallicula* (Freeman, 1967), *Mnemiopsis* (Henry and Martindale, 2000;Tamm, 2012b;Bading et al., 2017;Ramon-Mateu et al., 2019;Edgar et al., 2021;Ramon-Mateu et al., 2022;Mitchell et al., 2024), *Bolinopsis* (Moroz, 2024) and *Pleurobrachia* (Jager et al., 2008;Alié et al., 2011;Tamm, 2012a) suggest that several cell types, early ‘injury’ response transcription factors and a variety of signal molecules are recruited in ctenophores’ fusion processes.

Consequently, injury-related signaling (Moroz et al., 2021b;Moroz et al., 2023), mesoglean ameboid cells (Traylor-Knowles et al., 2019), and the widespread multifunctionality of macrophages are likely involved in the initial repair processes (Mitchell et al., 2024). The polarity of muscles (Scimone et al., 2017;Feige et al., 2018), as co-organizers of regeneration, should be considered.

In the quest for mechanisms of real-time physiological coordination, both neural and non-neuronal/immune integrative systems (Wenger et al., 2014;Traylor-Knowles et al., 2019)) should be equally studied. In addition to canonical electric and chemical synapses, we reported that two polygonal neural networks between two fused *Pleurobrachia* form morphological and functional connectivity, indicating that neural elements have rapidly regenerated at the injury site.

Burkhard and colleagues showed that in the cydippid larvae of *Mnemiopsis* 5 out of 33 studied neurons formed a syncytium with fused plasma membranes of neurites (Burkhardt et al., 2023). However, synaptic and volume transmission remains the dominant mode of intercellular signaling in ctenophores (Moroz, 2023). Both ancestral complements of small transmitters and secretory peptides (Moroz et al., 2021b) can form dynamic chemoconnectomics of fusion, with ‘collective intelligence of cells’ (Levin, 2023) (Moroz and Romanova, 2023) as the foundation for morphological homeostasis.

As perspective reductionistic paradigms, we emphasize the importance of semi-autonomous organoid-type entities, named here as neurobots or ctenobots. These were rapidly generated with an astonishing diversity of shapes and can be made from virtually all organs, starting from the aboral organ, combs, and auricles. Neurobots were formed within hours and could live over a week in seawater without any nutrients. In technical terms, neurobots/ctenobots are ‘synthetic living machines’ analogous to those fabricated from *Xenopus* as xenobots (Blackiston et al., 2021). This type of experimental bioengineering opens new perspectives and opportunities for biologically inspired robotics (Pfeifer et al., 2007) and deciphering mechanisms of systemic integration in free-behaving simplified paradigms.

We also think about the injury as a trigger and a powerhouse for the evolution of multicellularity via systemic stress responses (Love and Wagner, 2022). Here, the neurogenic role of injury (Moroz, 2009;2014) and “memory of injury (Walters and Moroz, 2009) can be exaptations of early metazoan innovations, including high regeneration potential (Hulett et al., 2024), and even facilitate making colonial organisms at the dawn of animal life.

### Conclusion

Ctenophores are ‘aliens of the sea’ with complex bilaterian-grade organization. They apparently independently evolved key animal traits such as neurons, muscles, mesoderm, and through-gut. As emerging reference species, sister to the rest of Metazoa, they deserve greater attention for integrative real-time physiological analyses at all levels of biological organization, from behaviors to cells, focusing on intracellular communications. Neither neural nor immune integrative mechanisms are currently known for ctenophores.

1. Ctenophore immune systems might also be unique, with a reduced capability for allorecognition. This enables a broad spectrum of grafting and fusion experiments resulting in chimeric animals, as documented for three species of ctenophores (*Pleurobrachia, Bolinopsis*, and *Mnemiopsis*).
2. Such broader taxonomic occurrences of grafting and fusion capabilities suggest that this trait might occur in the common ctenophore ancestor, although parallel origins of extensive regeneration potential across different ctenophore lineages can not be excluded and more comparative studies are required.
3. Ctenophore neural and immune systems can be coupled, and the polygonal neural networks establish morphological connections and functional continuity between two fused individuals (Fig. 4). The contribution of synaptic and non-synaptic mechanisms of cross-individual communications is currently unknown and can be resolved by combining ultrastructural and physiological studies.
4. The experimentally obtained diversity of semi-autonomous neurobots and chimeric animals from different ctenophore species illuminates prominent phenotypic plasticity, enabling the design and bioengineering of hybrid neural systems and even entire animals.

Future directions require the identification of self-recognition/adhesive molecules and signaling integrative mechanisms in ctenophores, where injury-induced secretory molecules (e.g., NO, Glutamate, ATP, small peptides - (Moroz and Kohn, 2011;Moroz et al., 2021a;Moroz et al., 2021b)) can act as the primary fusion signals. In this endeavor, a reduced complement of ctenophore immune systems might facilitate fusion – the process relevant to the origin and early evolution of metazoans, and multicellularity in general. Practical biomedical outreach is equally relevant to the evolutionary analysis of tumor formations (Kapsetaki et al., 2023) as potential sources of innovations.

In sum, the unification of biodiversity, genomics, cell biology, and neuroscience opens unprecedented opportunities for experimental synthetic biology in the XXI century, from the reconstruction of ancient neural systems to making chimeric neural circuits, new brains, and even multicellular organisms.

## Methods

Small adult *Bolinopsis* (2-3 cm length) and *Pleurobrachia bachei* (1-2 cm) were collected from the dock at Friday Harbor Laboratories (University of Washington, USA) and maintained in seawater tanks with running seawater. For fusion experiments, two *Bolinopsis* animals of similar size were transferred into a Sylgard-coated petri dish. Then, a part of the body wall of each animal (i.e., 1/4-1/3 of the size of the animals) was cut off from one side. The metal insect pins were used to hold the animals together, with cut surfaces facing each other. Then (after an hour, when animals started to fuse), the Petri dishes were transferred to a refrigerator (5° C) or seawater tank with running seawater (10-14° C) for 12-20 hours. The animals were fused entirely the next day and were transferred to a clean glass dish for taking pictures through a dissecting microscope using AmScope camera or iPhone. Similar experiments were performed on *Pleurobrachia* with the only difference in timeline – the whole regeneration process was significantly slower in this species by a factor of 2. Therefore, after day one, the fusion process started but was not completed, and animals were released in the seawater tank for one additional day for complete healing and fusion before final imaging.

To look at the tissue microstructure in ctenobots, we labeled the muscle fibers with phalloidin (Alexa Fluor 568 conjugated from Molecular Probes), which binds to F-actin. The ctenobots were fixed overnight in 4% paraformaldehyde in 0.1 M PBS (phosphate-buffered saline; pH=7.6) in a refrigerator for 12 hours and then washed for 8-12 hours using multiple PBS rinses. The samples were incubated in PBS with phalloidin at a final dilution of 1:80 for 12 hours and then washed in several PBS rinses for 6 hours.

To visualize the nervous system, we used anti-tubulin antibodies according to the following protocol (see details in (Norekian and Moroz, 2020;2024a). Briefly, after fixation and washing in PBS, the samples were incubated for 12 hours in a blocking solution of 6% goat serum in PBS and then for 48 hours in the primary anti-tubulin antibody (Bio-Rad, Catalog# MCA77G, RRID: AB_325003) diluted in blocking solution at a final dilution 1:50 in a refrigerator at 5ºC. Following a series of PBS washes for 12 hours, the samples were incubated for 24 hours in secondary goat anti-rat IgG antibodies (Alexa Fluor 488 conjugated; Molecular Probes, Invitrogen, WA, MA, Catalog# A11006, RRID: AB_141373) at a final dilution of 1:40 and then washed in a series of PBS rinses for 12 hours. To stain the nuclei, the preparations were mounted in Antifade Mounting Medium with DAPI (Vectashield; Cat#H-2000) on glass microscope slides. The slides were viewed and photographed using a Nikon Microscope Eclipse E800 using standard TRITC and FITC filters and Nikon C1 laser scanning confocal attachment.

## Acknowledgments

This research was supported by National Science Foundation grants (2341882) awarded to L.L.M. Additionally, this work was partly funded by the National Institute of Neurological Disorders and Stroke of the National Institutes of Health under Award Number R01NS114491 to L.L.M.

